# A connectivity-constrained computational account of topographic organization in primate high-level visual cortex

**DOI:** 10.1101/2021.05.29.446297

**Authors:** Nicholas M. Blauch, Marlene Behrmann, David C. Plaut

## Abstract

Inferotemporal cortex (IT) in humans and other primates is topographically organized, containing multiple hierarchically-organized areas selective for particular domains, such as faces and scenes. This organization is commonly viewed in terms of evolved domain-specific visual mechanisms. Here, we develop an alternative, domain-general and developmental account of IT cortical organization. The account is instantiated as an Interactive Topographic Network (ITN), a form of computational model in which a hierarchy of model IT areas, subject to connectivity-based constraints, learns high-level visual representations optimized for multiple domains. We find that minimizing a wiring cost on spatially organized feedforward and lateral connections within IT, combined with constraining the feedforward processing to be strictly excitatory, results in a hierarchical, topographic organization. This organization replicates a number of key properties of primate IT cortex, including the presence of domain-selective spatial clusters preferentially involved in the representation of faces, objects, and scenes, columnar responses across separate excitatory and inhibitory units, and generic spatial organization whereby the response correlation of pairs of units falls off with their distance. We thus argue that domain-selectivity is an emergent property of a visual system optimized to maximize behavioral performance while minimizing wiring costs.

**Significance Statement:** We introduce the Interactive Topographic Network, a framework for modeling high-level vision, to demonstrate in computational simulations that the spatial clustering of domains in late stages of the primate visual system may arise from the demands of visual recognition under the constraints of minimal wiring costs and excitatory between-area neuronal communication. The learned organization of the model is highly specialized but not fully modular, capturing many of the properties of organization in primates. Our work is significant for cognitive neuroscience, by providing a domain-general developmental account of topographic functional specialization, and for computational neuroscience, by demonstrating how well-known biological details can be successfully incorporated into neural network models in order to account for critical empirical findings.

---

Inferotemporal cortex (IT) subserves higher-order visual abilities in primates, including the visual recognition of objects and faces. By adulthood in humans, IT cortex, and ventral temporal cortex more generally, contains substantial functional topographic organization, including the presence of domain-selective spatial clusters in reliable spatial locations, including clusters for faces (1–3), objects (4), buildings and scenes (5, 6), and words (7). Similar domain-level topographic properties have been found in rhesus macaque monkeys, including multiple regions of clustered face selectivity (8–10). Intriguingly, this selectivity is encompassed in a larger scale “mosaic” of category-selectivity, in which areas of category-selectivity themselves have further columnar clustering within them (11–13), pointing to more general principles of organization beyond the domain level. In line with this idea, human IT cortex also exhibits larger-scale organization for properties such as animacy and real-world size (14, 15), and midlevel features characteristic of these properties and domains have been shown to account well for patterns of high-level visual selectivity (16). How these domain-level and more general facets of functional organization arise, how they are related, and whether and in what ways they rely on innate specification and/or experience-based developmental processes remain contentious.

Recent work has demonstrated that the neural basis of face recognition depends crucially on experience, given that deprivation of face viewing in juvenile macaque monkeys prevents the emergence of face-selective regions (17). Relatedly, the absence of exposure to written forms through reading acquisition precludes the emergence of word-selective regions (18, 19). That there exists clustered neural response selectivity for evolutionarily new visual categories such as written words offers further evidence that the topographic development of the human visual system has a critical experience-dependent component (20, 21). In contrast with a system in which innate mechanisms are determined through natural selection, this experiential plasticity permits the tuning of the visual system based on the most frequent and important visual stimuli that are actually encountered, thereby enabling greater flexibility for ongoing adaptation across the lifespan.

There is considerable computational evidence that experience-dependent neural plasticity can account for the response properties of the visual system at the single neuron level. Classic work demonstrated that the statistics of natural images are sufficient for learning V1-like localized edge-tuning within a sparse coding framework (22, 23). More recently, deep convolutional neural networks (DCNNs) trained on image classification have been successful in accounting for the tuning of neurons in V1, V2, V4, and IT in a hierarchically consistent manner, where deeper layers of the DCNN map onto later layers of the anatomical hierarchy (24, 25).

Above the single-neuron level, considerable prior work has demonstrated that topographic organization in V1 may emerge from self-organizing, input-driven mechanisms (26–32) (for review, see 33). For example, the pinwheel architecture of spatially repeating smooth orientation selectivity overlaid with global retinotopy has been shown to be well-accounted for by Self-Organizing Maps (SOMs) (29, 30, 34). One notable application of an SOM to modeling high-level visual cortex by Cowell and Cottrell (35) demonstrated stronger topographic clustering for faces compared to other object categories (e.g., chairs, shoes), suggesting that the greater topographic clustering of faces in IT is due to greater within-category similarity among faces compared to these other categories. This work provides a strong case for domain-general developmental principles underlying cortical topography in IT, but at least two important issues remain unaddressed. First, rather than only supporting discrimination of face from non-face categories (as in 35), face representations in humans (and likely non-human primates, though see (36)) must support the more difficult and fine-grained task of individuation; this task requires a “spreading transformation” of representations for different face identities (37, 38), which could could alter the feature space and its topographic mapping, and necessitate a more domain-specialized representation than arises in an SOM. And secondly, rather than a single face-selective area, IT cortex actually contains multiple hierarchically-organized face-selective regions with preferential inter-connectivity (39). Generally, SOMs are not well equipped to explain such hierarchical topographic interactions, as they are designed to map a feature space into a topographic embedding, but not to transform the feature space hierarchically in the way needed to untangle invariant visual object representation from the statistics of natural images (40). This suggests that SOMs may not be a good model of topographic development in cortical networks.

An alternative approach to studying topographic organization involves incorporating distance-dependent constraints on neural computation within more general neural network models (41–44). Of particular interest is a hierarchical neural network developed by Jacobs and Jordan (43) in which error-driven learning was augmented with a spatial loss function penalizing large weights to a greater degree on longer versus shorter connections. This model was shown to develop topographic organization for ‘what’ versus ‘where’ information when trained with spatially segregated output units for the two tasks. Closely related work by Plaut and Behrmann (45) demonstrated that a similar spatially-constrained model with biased demands on input (e.g., retinotopy) and output (e.g. left-lateralized language) could account for the organization of domain-specific areas in IT cortex, such as the foveal bias for words and faces, leftward lateralization of words, and rightward lateralization of faces (46–48). However, to date, none of these structurally-biased neural network models have been applied to large-scale sets of naturalistic images, the statistics of which are thought to organize high-level visual representations in IT cortex (49), and the topography in these models (43, 45) has been analyzed at a relatively coarse level. Nonetheless, this early work raises the possibility that the application of distance-dependent constraints in a modern deep neural architecture trained on natural images might provide a more comprehensive account of topographic organization in IT.

Recently, Lee and colleagues (50) have modeled the topography of IT cortex with a deep neural network trained on a large set of natural images, using a correlation-based layout that explicitly encouraged units within a layer of the network to be spatially nearer to units with correlated responses, and farther from units with uncorrelated or anti-correlated responses. As a result, the network developed face-selective topography that corresponded well with data from macaque monkeys. However, this approach *imposes* topographic functional organization on the network based on measured functional responses, rather than *deriving* it from realistic principles of cortical structure and function, such as constraints on connectivity. Moreover, like the SOM, the approach can explain only *within-area* topographic organization, and not relationships between areas, such as multiple stages of IT cortex and their interactions with upstream and downstream cortical areas. Thus, the question remains whether such basic structural principles can account for the topographic organization of IT.

In the current work, we combined the approaches of task-optimized DCNN modeling (49, 50) with flexible connectivity-constrained architectures (43, 45) to develop a hierarchical model of topographic organization in IT cortex. We implemented a bias towards local connectivity through minimization of an explicit wiring cost function (43) alongside a task performance cost function. Intriguingly, we observed that this pressure on local connectivity was, on its own, insufficient to drive topographic organization in our model. This led us to explore two neurobiological constraints on the sign of connectivity—strictly excitatory feedforward connectivity, and the separation of excitation and inhibition—with the result that both, and particularly excitatory feedforward connectivity, provided a powerful further inductive bias for developing topographic organization when combined with a bias towards local connectivity.

## Results

### A connectivity-constrained model of ventral temporal cortex produces hierarchical, domain-selective response topography

Our Interactive Topographic Network (ITN) framework for modeling high-level visual cortex consists of an *encoder* that approximates early visual cortex, followed by *interactive topography* areas that approximate IT cortex (Figure 1A; see Methods for details). We first present the results of simulations of a specific ITN model, in which a ResNet-50 encoder is pretrained on a large dataset including several categories from the domains of objects, faces, and scenes (each domain matched in total training images). The trained encoder provides input to a 3-area IT with separate posterior (pIT), central (cIT), and anterior (aIT) areas. Each IT area consists of separate banks of excitatory (E) and inhibitory (I) units, and feedforward connectivity between areas is limited to the E units. After training, the model performed well on each domain, reaching a classification accuracy of 86.4% on the face domain, 81.8% on the object domain, and 65.9% on the scene domain (see Supplementary Figure S1). Performance differences across domains are unlikely to be an artifact of the specific architecture as they can be seen across a variety of CNNs, reflecting the intrinsic difficulty of each task given the variability within and between categories of each domain for the given image sets.

**Fig. 1.**
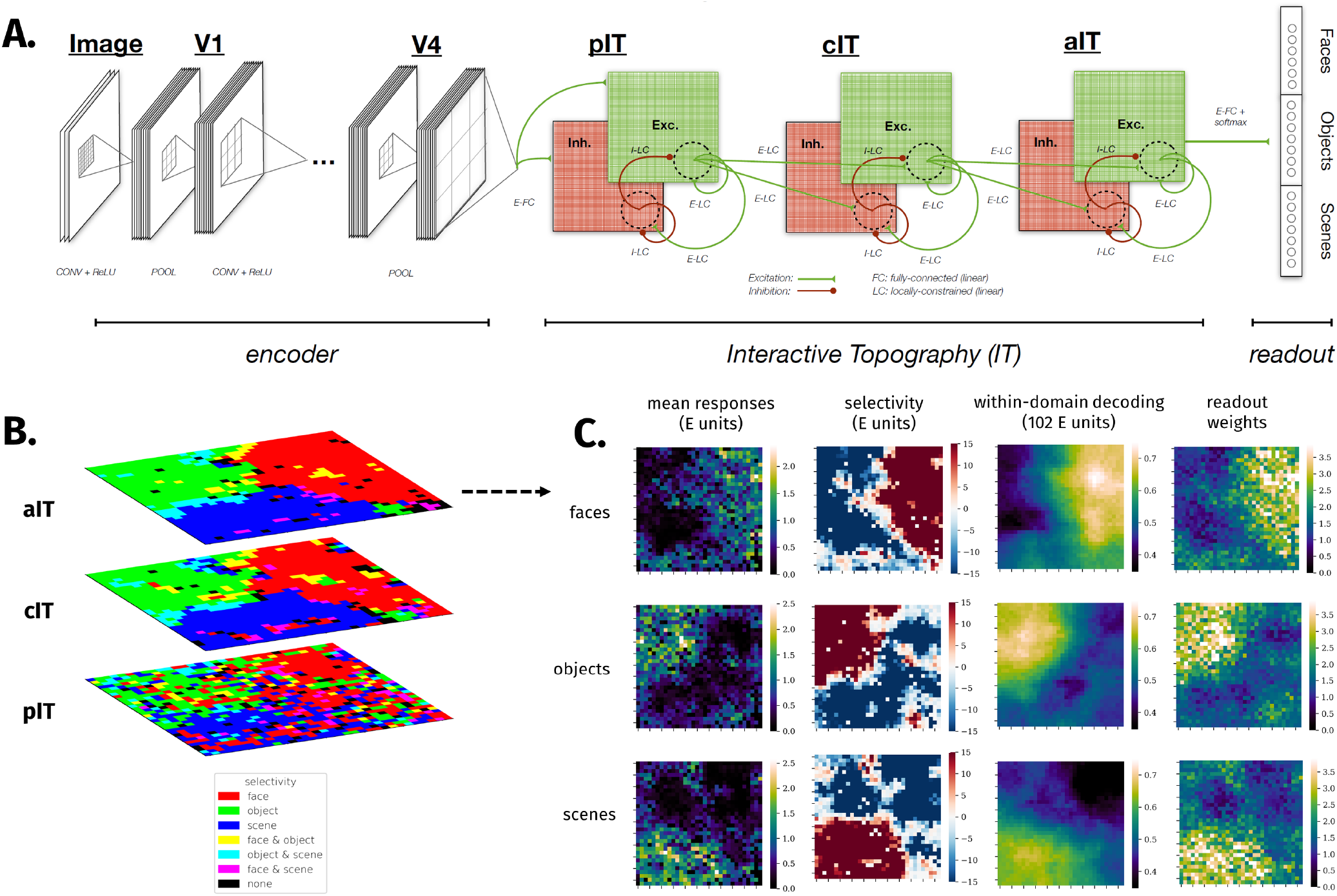
The Interactive Topographic Network produces hierarchical domain-level organization. **A.** diagram of the Interactive Topographic Network (ITN). An ITN model consists of three components: an *encoder* that approximates early visual processing prior to inferotemporal cortex, the *interactive topography* (IT) areas that approximate inferotemporal cortex, and the *readout* mechanism for tasks such as object, scene, and face recognition. The architecture of each component is flexible. For example, a 4-layer simple convolutional network or a deep 50-layer ResNet can be used as the encoder; whereas the former facilitates end-to-end training along with a temporally-precise IT model, the latter supports better learning of the features that discriminate among trained categories. In this work, topographic organization is restricted to the IT layers. The figure depicts the main version of the ITN containing three constraints: a spatial connectivity cost pressuring local connectivity, separation of neurons with excitatory and inhibitory influences, and the restriction that all between-area connections are sent by the excitatory neurons. The final IT layer projects to the category readout layer containing one localist unit per learned category, here shown organized into three learned domains. (Note that this organization is merely visual and does not indicate any architectural segregation in the model. **B.** Domain selectivity at each level of the IT hierarchy. Selectivity is computed separately for each domain, and then binarized by including all units corresponding to *p <* 0.001. Each domain is assigned a color channel in order to plot all selectivities simultaneously. Note that a unit can have zero, one, or two selective domains, but not three, as indicated in the color legend. **C.** Detailed investigation of domain-level topography in aIT. Each heatmap plots a metric for each unit in aIT. The first column shows the mean domain response for each domain, the second column shows domain selectivity, the third column shows the within-domain searchlight decoding accuracy, and the fourth column shows the mean of weights of a given aIT unit into the readout categories of a given domain.

The trained model exhibits domain-level topographic organization that is hierarchically linked across corresponding sectors of each layer (see Figure 1B). This result reflects the fact that the distance-dependent constraints on feedforward connectivity pressured units that have minimal between-area distances to learn a similar tuning, which means that each layer is roughly overlapping in their respective (separate) 2D topography. The topographic organization gets somewhat smoother moving from pIT to cIT, most likely because units in cIT and aIT (but not pIT) have local feedforward receptive fields and thus greater constraint on local cooperation.

We next scrutinized the topography in aIT, where there are very smooth domain-level responses, and where we can directly compare responses with those of the recognition readout mechanism. We computed mean domain responses, plotted in the first column of Figure 1C, and domain selectivity, plotted in the second column, which demonstrates corresponding topographic organization. We confirmed the functional significance of response topography by conducting a searchlight analysis inspired by multivariate approaches to analyzing functional magnetic resonance imaging (fMRI) data (51). We used searchlights containing the 10% (102) nearest units. The results of this analysis, shown in the third column of Figure 1C, revealed topographic organization of information for discriminating between categories of each domain that is strongly correlated with the domain selectivity maps for each domain (all *ps <* 0.0001).

To further confirm the functional significance of the topographic organization, we analyzed the spatial organization of readout weights from aIT to the localist category readout layer. We evaluated whether each domain placed more weight in reading out from the units for which there was greater selectivity, by calculating the mean domain response weight for each unit, averaged over classes in each domain. This produced a map for each domain, shown in the last column of Figure 1C. We find a large positive correlation between the mean readout weight and the mean response for each domain (all *r*s>0.7, all *p*s*<* 0.0001), further demonstrating the functional significance of the response topography.

### Excitatory and inhibitory units operate as functional columns

Thus far we have focused on the representations in the E cells, both for convenience and clarity, and because it is the E units that exclusively project to downstream areas (including the category readout units). We next assessed whether the I units show a similar topographic organization, and whether it is linked with the E cells. The selectivity for E and I cells is plotted and correlated in Figure 2. The I cells show similar domain-selective topography to the E cells. Moreover, the activities of E and I units in the same 2D location have highly correlated activities over each domain of images, as well as over all images. If we consider a pair of E and I neurons at a given location on the 2D sheet to correspond to a cortical column, our result is reminiscent of the finding that biological neurons in different layers at the same location on the 2D flattened cortex have similar response properties (52). In this way, E and I units in the model appear to act as functional columns.

**Fig. 2.**
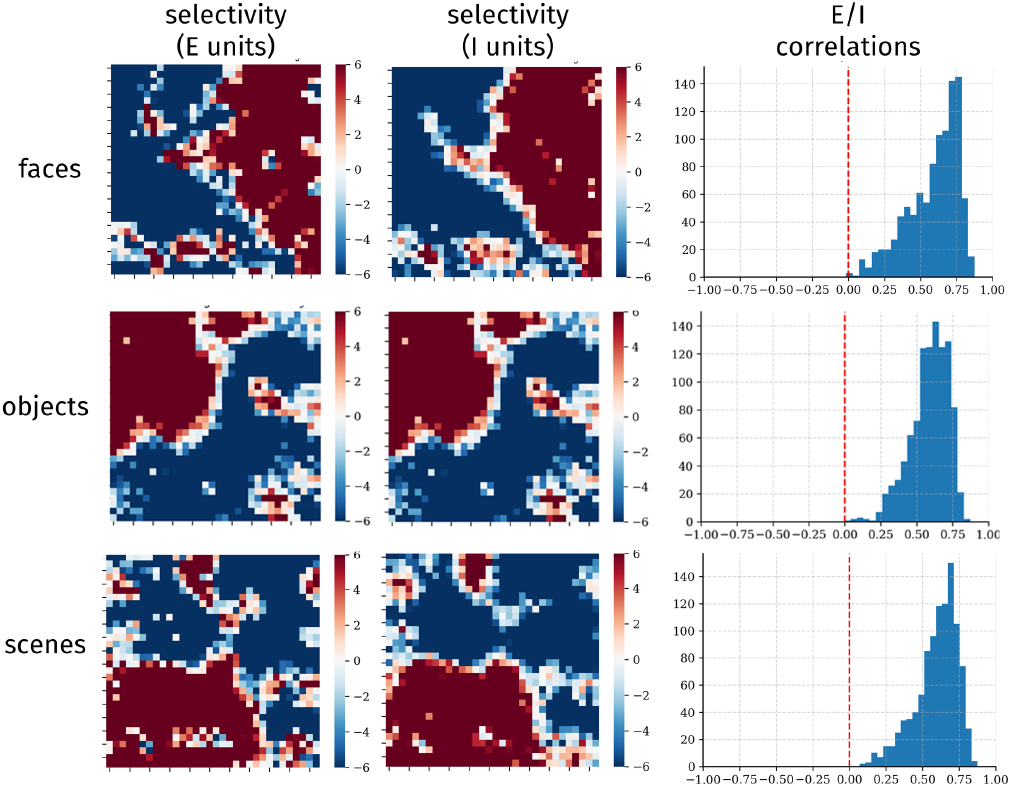
E and I cells act as functional columns. Selectivity of cIT excitatory (E) units (left columns), and inhibitory (I) units (middle column) for each domain, and histograms of response correlations between co-localized E and I units for images from each domain (right column).

### Effects of lesions indicate strong yet graded domain-level specialization

We next performed a series of “lesion” analyses in the model in order to compare with neuropsychological data on face and object recognition (53–55). First, we performed focal lesions, as would be experienced by most patients with acquired brain damage. To simulate the impairment of patients with maximally specific deficits, we centered circular focal lesions of various sizes at the center of (smoothed) domain selectivity. Performance following each lesion was measured separately for each domain.

The results of this lesion analysis are shown in Figure 3A. Focal lesions centered on each domain for two representative lesion sizes—using 20% and 30% of the aIT units—are shown in Figure 3A. Focal lesions centered on each domain lead to an especially severe deficit in recognition for that domain, and milder but significant deficits for the other domains as well. For a medium sized lesion of 20% of the units (Figure 3A, right), the deficit is significant for all domains (all *p*s<0.05), and significantly stronger for recognition of the target domain (all *p*s<0.05).

**Fig. 3.**
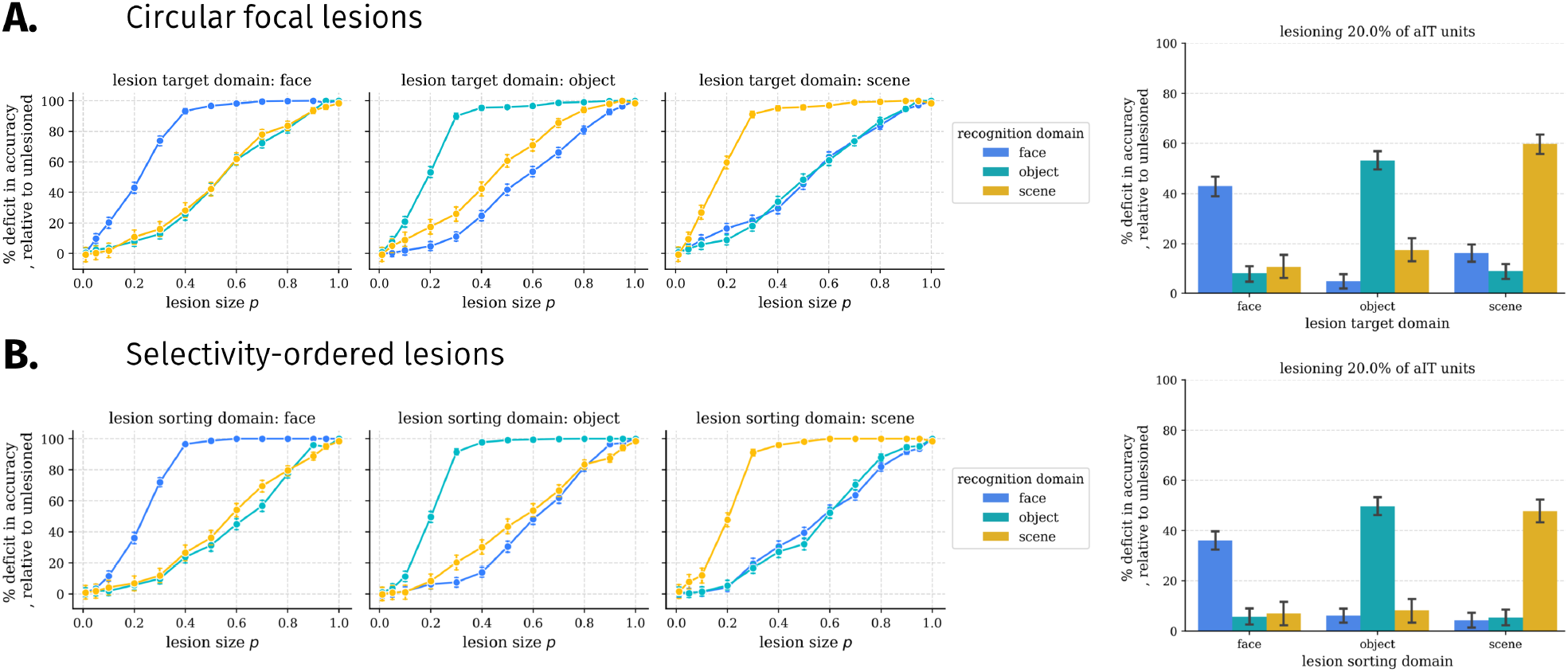
Lesion results in the ITN model. Each plot shows the relative effects of a set of same-sized lesions on recognition performance for each domain, relative to the performance on the same domain in the undamaged model. Error bars show bootstrapped 95% confidence intervals over trials; thus, the statistical significance of a given lesion can be assessed by determining whether the confidence interval includes 0. **A.** Damage from circular focal lesions centered on the peak of smoothed selectivity for each domain. Left: results for a variety of lesion sizes. Right: a focused analysis of an intermediate lesion size of 20% of the aIT units. **B.** Damage from selectivity-ordered lesions for each domain. Left: results for a variety of lesion sizes. Right: a focused analysis of an intermediate lesion size of 20% of the aIT units.

Are these more general effects of circumscribed lesions on non-preferred domains the result of imperfect (patchy) or non-circular topographic organization of an underlying modular organization? To answer this question, we performed selectivity-ordered lesions, in which units were sorted by their selectivity for a given domain, and selected according to their sorting index, shown in Figure 3B. The effects of damage in this case are similar to those for focal lesions, with greater damage to the domain on which sorting was performed, and smaller deficits to other domains for lesions targeting at least 20% of the units. Specifically, for 20% lesions, we found smaller but still significant deficits for both the preferred and non-preferred domains compared to focal lesions. This suggests that some but not all of the damage to the non-preferred domain induced by focal lesions may be due to imperfect or non-circular topographic functional organization. Importantly, these more distributed effects of lesions indicate that the functional organization, while highly specialized, is not strictly modular, at least with respect to one influential definition of modularity (56). Supplementary Figures S3, and S4 provide additional data on the nature of domain specialization in the network.

### Domain selectivity exists within a broader organization similar to that of primate IT cortex

Previous empirical research has demonstrated that the response correlations between pairs of neurons fall off smoothly with increasing distance between the neurons (data from 57, as plotted in (50), Figure 4A.). This finding has been used to develop a class of topographic neural network models that explicitly fits the spatial layout of units to this relationship (50). We explored whether this relationship emerged naturally in our network due to its constrained connectivity, in line with the emergence of domain-selective topography. We thus computed the correlations among pairs of unit activations across images as a function of the distance between the units, focusing on aIT. As shown in Figure 4B, there is, indeed, a smooth decay of response correlations with distance, matching the qualitative trend in the empirical data (50, 57).

**Fig. 4.**
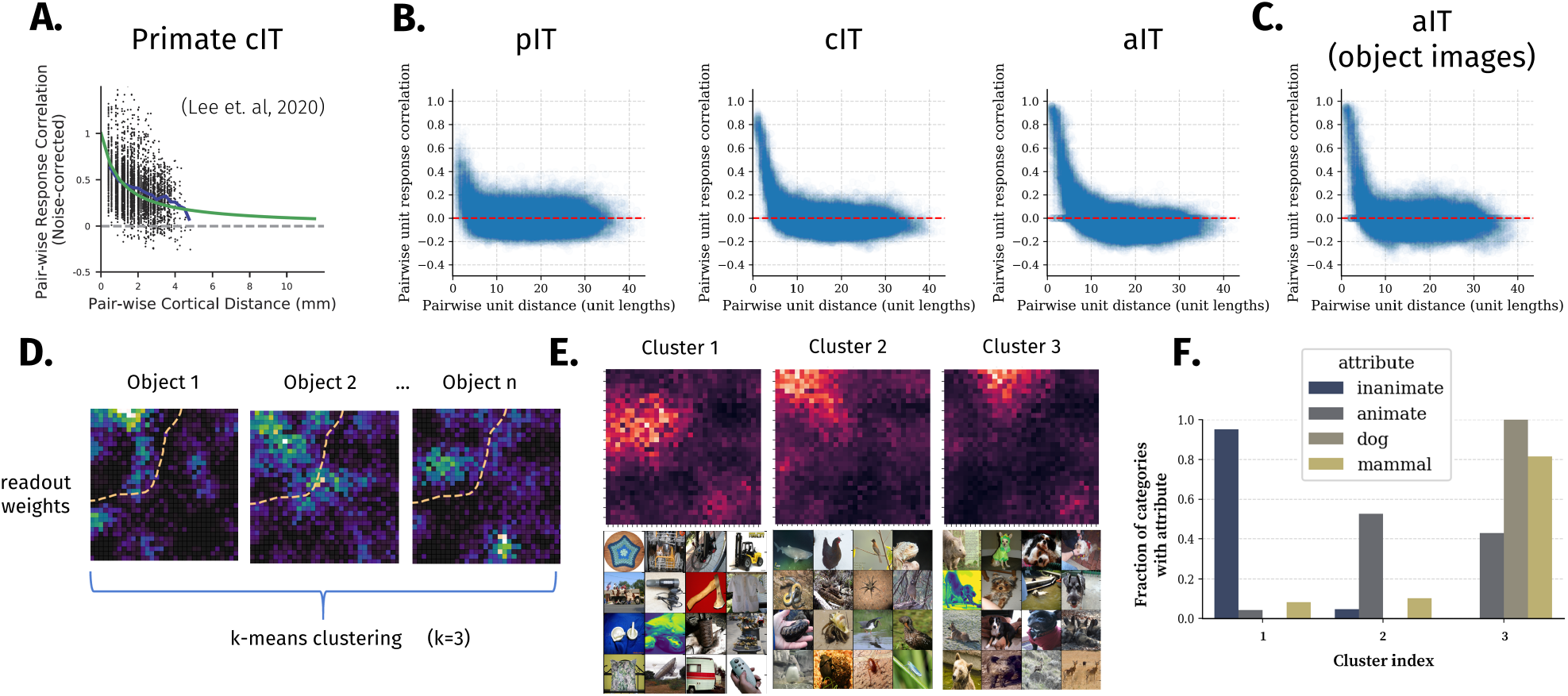
Generic topographic organization beyond domain-selectivity emerges through task optimization under biologically-plausible constraints on connectivity. **A.** Distance-dependent response correlation in macaque IT (reproduced from 50, per CC-BY-NC-ND license). **B.** Distance-dependent response correlation in the excitatory cells of each layer, using images from all three domains (objects, faces, scenes). **C.** Distance-dependent response correlation in aIT using images from the object domain only, highlighting within-domain generic functional organization. **D.** Schematic for within-domain readout weight clustering analysis. The readout weights for each category of a given domain (i.e. objects) are subject to a k-means clustering analysis. *k* = 3 clusters are used to identify dominant patterns of variation across categories in the information read-out by downstream localist category units. The output of the analysis is a cluster centroid and set of cluster category members for each of *k* clusters. **E.** Results of the within-domain readout weight clustering analysis. Top row: cluster centroids; bottom row: sample of 16 cluster category members; the dashed box at top-right re-plots the aIT domain-level selectivity for comparison with the object within-domain topography. **F.** Attribute quantification of category membership in the readout weight cluster analysis. For each cluster and attribute, the fraction of total categories with the attribute that are present in the cluster is plotted.

This result is not simply due to differences between domains, as it is also found when examining responses to images within each domain separately (shown for objects in Figure 4C). Along with previous results (50), our findings suggest that the domain-level topography may simply be a large-scale manifestation of a more general representational topography in which the information represented by neighboring units is more similar than that represented by more distal units. Importantly, our results go beyond previous ones to also demonstrate that this organization can arise under explicit wiring length and sign-based constraints on connectivity.

The generic distance-dependent functional relationship just discussed would suggest that functional organization may be exhibited at finer scales than the domain level. To assess this, we performed a clustering analysis on the readout weights from aIT. We adopted this approach due to the similarity between the readout weights and response topography in aIT (Figure 1C). A given category will achieve its maximal output response when the activation pattern in aIT most closely aligns with the readout weights. Thus, the readout weights for a given category act as a sort of category template to match with representations in aIT. Clustering the readout weights directly, rather than interpreting a set of activations to natural images, enables clustering solutions to be explicitly linked to each category. This allows for a concise clustering solution containing one element for each category: the readout weights projecting from aIT to the identity unit for that category. We thus performed k-means clustering on the readout weights of all categories separately for each domain using *k* = 3 clusters (Figure 4D), finding the centroids of these clusters, and visualizing them in the 2D layout of aIT. The centroids and cluster category members are shown in Figure 4E. The cluster centroids show smooth topographic organization, with each cluster having a primary hot-spot of weight, and graded weight in other parts of aIT. Visual inspection of the cluster category members suggests a striking organization for different classes of object categories. This organization is confirmed through cluster assignment quantification in Figure 4F. The first two clusters represent the vast majority of animate categories, with the first cluster representing mostly non-mammalian animate categories such as birds and reptiles, and the second cluster representing mostly dogs and other mammals such as bears and raccoons. Last, the third cluster represents the vast majority of inanimate objects such as clocks and various tools. Further analysis of the scene and face domain readout weights indicated a similar within-domain organization, with scenes being clustered by indoors-outdoors and natural-manmade dimensions, and faces being clustered by gender and hair color dimensions (Supplementary Figures S7, S8).

### Networks can reduce spatial costs and maintain performance by increasing topographic organization

The optimization problem introduced by Jacobs and Jordan (43) and employed in this work (Equation 4) explicitly works to both maximize visual recognition performance through a task-based loss term ℒ_*t*_, and to minimize overall wiring cost through a connection-based lost term ℒ_*w*_ that scales with the square of connection distance. To what extent does minimizing this wiring cost term compromise performance? To answer this question, we tested multiple ITN models with varying wiring cost penalties *λ*_*w*_ and measured the resulting wiring cost and task performance. We computed wiring cost in two ways. The first way is by using the ℒ_*w*_ term, which takes into account both the length and strength of connections. The second way is inspired by the wiring cost minimization framework (58), which cares only about the presence—rather than the strength—of connections, along with their distance. To compute this wiring cost ℒ_*w,u*_, we sparsified the network to contain only the 1% strongest connections (sparsity=0.99), and took the averaged squared distance of remaining connections (59, see Equation 6); this sparsification introduces minimal performance deficits in the main ITN model (and Figure 5A). The results, shown in Figure 5A., demonstrate that increasing the wiring cost penalty *λ*_*w*_ by an order of magnitude decreased the first spatial cost ℒ_*w*_ by roughly an order of magnitude. Precisely, the log-log plot in Figure 5A (left) revealed a power law relationship of the form *y* = *Ax*^*m*^, where *m* = −1.24 (*p <* 0.001). The unweighted wiring cost *L*_*w,u*_ similarly decays roughly linearly on the log-log plot up to *λ*_*w*_ = 0.1, after which *L*_*w,u*_ saturates and then rises for increasing values of *λ*_*w*_. Thus, an intermediate value of *λ*_*w*_ appears sufficient to drive the network towards preferentially local connectivity, and further increasing *λ*_*w*_ may minimize further the optimization term ℒ_*w*_ through other means, such as by further shrinking small long-range weights and reducing participation at the grid boundaries where mean connection lengths are longest (see Figure 5C, top right). In contrast to the wiring costs, the final classification performance was only marginally affected by *λ*_*w*_ (log-log slope *m* = −0.0016, *p <* 0.001, explained variance *r*^2^ = 0.582; fit was not significantly better than log-linear regression, *m* = −0.0028, *p <* 0.001, explained variance *r*^2^ = 0.583) and the final top5 classification performance was unaffected by *λ*_*w*_ (*p >* .1; see Figure 5B). Last, increasing the wiring cost penalty gradually resulted in the emergence of domain-selective areas, along with distance-dependent pairwise response correlations (see Figure 5C). Thus, models with a large wiring cost penalty perform similarly to models with unconstrained connectivity but achieve very small wiring cost, through the development of topographic functional organization.

**Fig. 5.**
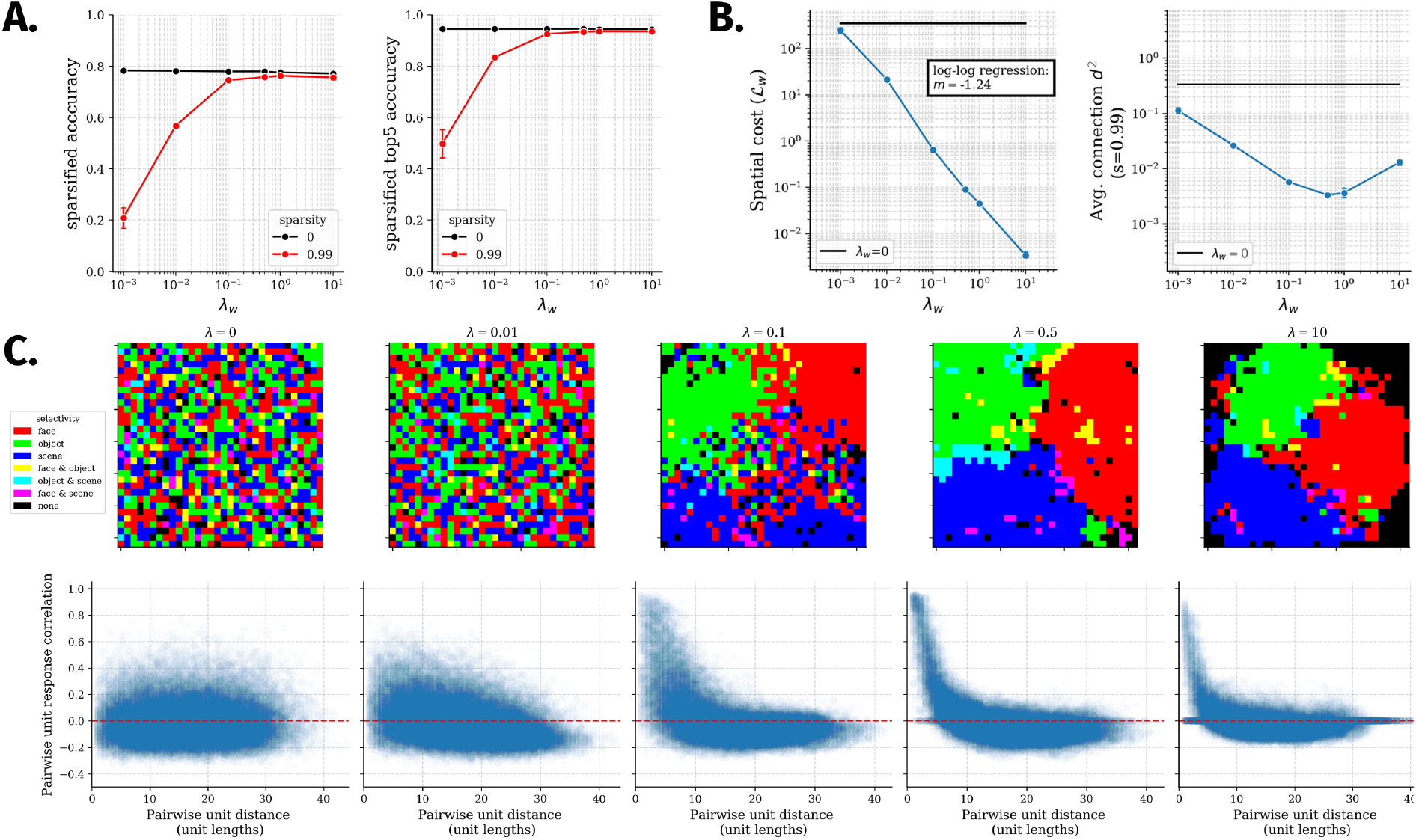
Spatial cost, visual performance, and emergent topography as a function of spatial wiring penalty *λ*_*w*_. Figures in **A-B** use 4 randomly initialized model instances per regularization strength. Error bars around markers show 95% confidence intervals for the plotted metric at a given spatial regularization strength, and black bands show 95% confidence intervals of the metric for a matched model without a spatial wiring penalty (*λ*_*w*_ = 0). **A.** Accuracy analysis. Left: mean top1 accuracy on validation images from all domains versus versus spatial wiring penalty *λ*_*w*_ on a log X-axis and linear Y-axis. Right: mean top5 accuracy plotted in the same manner. In both cases, results for the main model as well as a sparsified version of the model (fraction of *s* = 0.99 smallest within-IT weights set to 0) are plotted. **B.** Wiring cost analysis. Left: weighted spatial cost (*L*_*w*_) versus spatial wiring penalty *λ*_*w*_, plotted on log-X and log-Y axes. Right: unweighted spatial cost following sparsification versus spatial wiring penalty *λ*_*w*_, plotted on log-X and log-Y axes. **C.** Emergent topographic organization in one model instance of each spatial wiring penalty *λ*_*w*_. Top row: domain-selective aIT topography. Bottom row: generic aIT topography.

### Sign-based constraints are necessary for the development of topography

Having established that the main ITN architecture produces a variety of interesting and empirically grounded topographic organizational phenomena, we next performed a constraint-removal analysis to determine which constraints—in addition to the bias towards local connectivity—are necessary for the development of topographic organization. We varied three constraints: whether between-area feedforward connections were excitatory only, whether the model employed separate E and I unit populations within each area, and whether the model contained lateral (recurrent) connections within each area. We thus constructed four simplified models, comparing both domain-selective and generic topography with the full model used in earlier analyses.

The first reduced model, shown in the second column of Figure 6A, contained a bias towards local connectivity and recurrent connections but no sign constraints. This model did not develop domain-level topography, and yielded a very weak relationship of pairwise unit response correlation with distance. This indicates that the sign-based constraints were important for the development of topography in the main model. We next examined a model without the restriction that feedforward connections be limited to the excitatory neurons, but with separate neurons responsible for excitatory and inhibitory influences. Results for this model (Figure 6A, 3rd column) indicate that the E/I separation increased topography compared to the model without sign-based constraints. This model yielded a strong generic topographic organization, but a weaker domain-level topographic organization than the full model. We next examined a model without separate neurons for excitation and inhibition, but with the restriction that feedforward connections be excitatory. This model yielded strong domain-level and generic topographic organization. Lastly, we constructed a simple feedforward model in which we removed learned lateral connectivity, leaving only layer normalization to mediate within-area interactions. Like the previous model, this model yielded strong topographic organization both at the domain- and generic-levels.

**Fig. 6.**
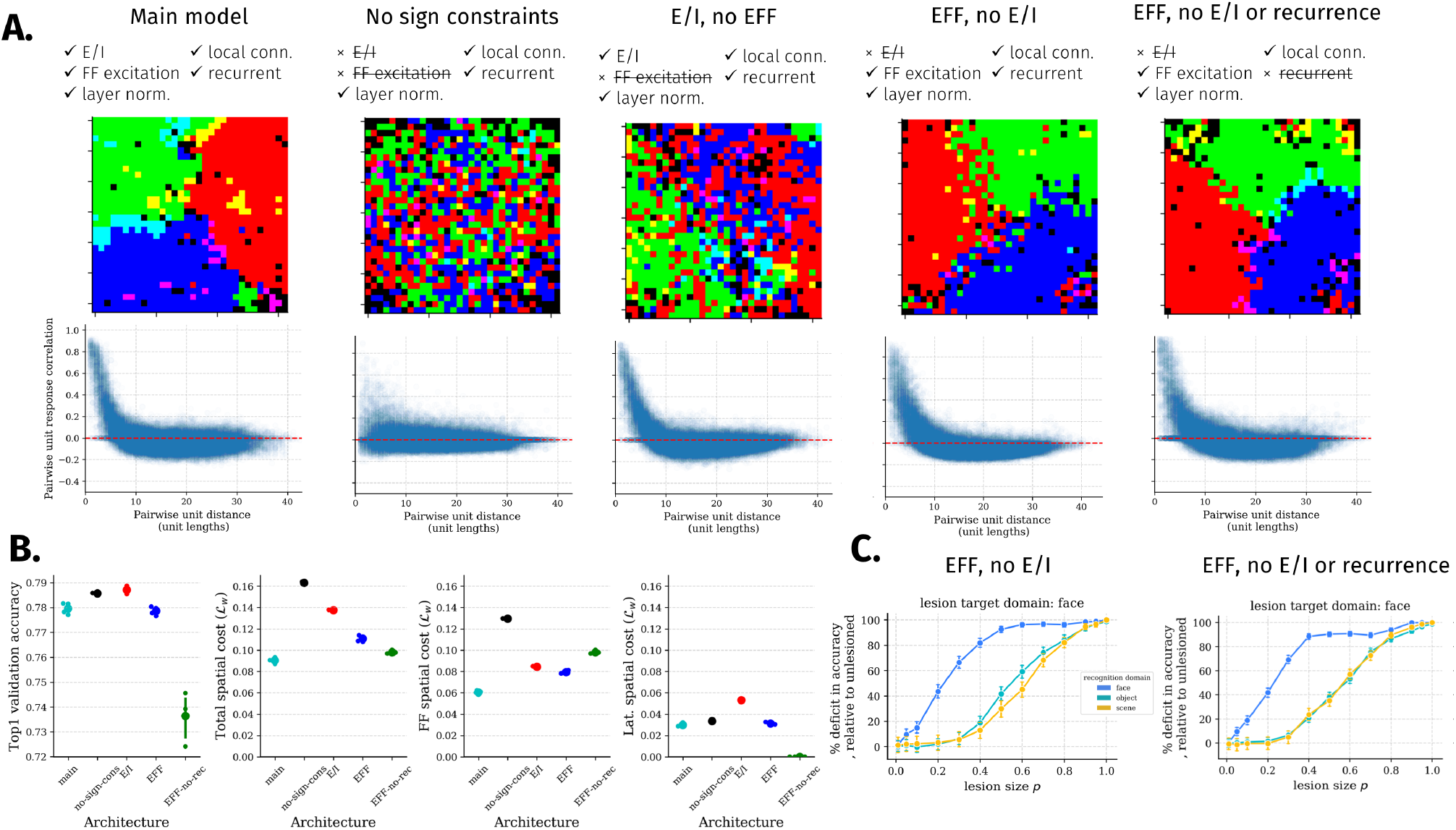
Constraint removal analyses. **A.** Domain-level and generic topography in layer aIT across models with different constraints implemented. From left to right: “main model” is the full model with separate excitation and inhibition, excitatory feedforward connections, learned recurrent connectivity, and a bias towards local connectivity; “no sign constraints” removes both sign-based constraints; “E/I, no EFF” has separate excitation and inhibition but both cell types send feedforward connections to the next layer; “EFF, no E/I” does not separate excitation and inhibition (one map of units), but feedforward connections are restricted to be excitatory; “EFF, no E/I or recurrence” does not separate excitation and inhibition, does limit feedforward connections to be excitatory, and does not have recurrent connections besides the layer normalization also implemented in all of the other models. **B.** Comparing performance and spatial cost of the main model with the variants shown in A. Left: final classification accuracy of 4 randomly initialized versions of each architecture. Second: final spatial cost. Third: final feedforward spatial cost, computed as the spatial cost of all between-area (feedforward) connections. Right: final lateral spatial cost, computed as the spatial cost of all within-area (lateral) connections. **C.** Domain-level functional specialization of a single model of each variant measured using face selectivity-ordered lesions. More complete results for these two models can be found in Supplementary Figures S5 and S6.

We next compared each of these model variants in their accuracy and spatial costs. First, we found that the accuracy of the recurrent models was very similar, with a very small (<1% point) advantage for models in which feedforward connectivity was not constrained to be excitatory. In contrast, accuracy for the feedforward model was reduced more substantially (>4% points), pointing to a performance benefit of the recurrent connections. Moreover, we found that, for the same *λ*_*w*_ across variants, the variants that developed clear domain-level organization had the smallest wiring cost (Figure 6B, 2nd panel). The variant without sign-based constraints— that demonstrated the least emergent topography—also had the highest wiring cost, and this was due to increases in feed-forward spatial costs (Figure 6B, 3rd panel). This can be understood in terms of this model requiring more weight over longer range connections due to the less ordered topography. Lastly, in addition to reduced performance, the feedforward variant yielded higher between-area spatial costs than the other topographically organized variants; these between-area spatial costs may be substantially more biologically burdensome than within-area spatial costs, since they incorporate between-area distances in addition to aligned point-to-point distances. For simplicity, and because modeling the complexities of cortical between-versus within-area distances was beyond the scope of this work, feedforward spatial costs in these models only include the aligned point-to-point distances. However, our results suggest that the feedforward model would have a difficult time reducing such costs, whereas the recurrent variants are able to minimize feedforward spatial costs through less expensive lateral connections.

Finally, we assessed the domain-specificity of the final two variants through the selectivity-ordered lesion approach used earlier (Figure 3). The results for face-selectivity-ordered lesions, shown in Figure 6C, indicate that the networks exhibit strong but graded specialization as in the main model, with somewhat weaker deficits at the small lesion size of 0.2 indicating somewhat stronger specialization. However, the rising deficits for all domains when lesion size is increased from 0.2 to 0.3, and from 0.3 to 0.4, strongly suggest that there is a partial graded overlap in the representation of domains, rather than a truly modular representation.

## Discussion

Is IT cortex a collection of independent, possibly hard-wired domain-specific modules, or a more general-purpose, interactive, and plastic system? The investigations presented here demonstrate that many of the key findings thought to support a modular view of separable, innately-specified mechanisms for the recognition of different high-level domains (faces, objects, scenes) can be accounted for within a learning-based account operating under generic connectivity constraints (also see 21). By simulating a biologically plausible Interactive Topographic Network (ITN) model of IT without domain-specific innate structure, we found that we can “let the structure emerge” (60, 61). Specifically, we observed that the model developed strongly domain-selective spatial clusters which contain preferential information for each domain, and which, when lesioned, produced largely (but not purely) specific deficits.

Beyond domain-level spatially clustered organization, the model exhibited a more generic form of topographic organization, whereby nearby units had more correlated responses over images compared to more distant units, a relationship which has been demonstrated in macaque IT cortex (50, 57). In combination with other modeling work (50) that pressured neurons to obey this relationship as a proxy “wiring” loss to develop face-selective topography, our work suggests that this generic spatial functional relationship appears to both underly domain-level organization and emerge from wiring cost minimization.

Importantly, wiring cost and task optimization (i.e., object, face, and scene image recognition), by themselves, were not sufficient to produce topographic organization: we found that two well-known biological details—excitatory-only between-area communication, and separate excitatory and inhibitory neural populations—could induce topographic organization in the context of wiring cost and task optimization. In particular, locally-biased excitatory feedforward connectivity provides an inductive bias that neighboring units should have positively correlated response properties, without specifying how correlated they should be. Since the network is constrained to perform multiple tasks, all units cannot be positively correlated to reach high performance; the network thus is encouraged to learn in a fashion whereby local units learn correlated representations and more distant units learn uncorrelated or anti-correlated representations, a hallmark of topographic organization (50). Additionally, the separation of excitation and inhibition contributed to topographic organization, but less so than the excitatory restriction on feedforward connectivity. We reason that the separation of excitation and inhibition serves to enhance topographic organization by inducing greater pressure on the lateral connections, since learned inhibitory responses must be mediated through lateral connections to and from the I cells. As this connectivity is biased to be local, it creates a pressure for local communication to be functionally smooth so that neurons representing related information can communicate with each other. While sign-based constraints played an important role in the development of topographic organization in this work, future work examining other tasks (62, 63) and architectures (64, 65) that place greater demands on lateral connectivity may find that local connectivity constraints suffice.

Our constraint-removal analysis allowed us to discover a simple model capable of producing many of the hallmarks of topographic organization in the main model. This feedforward variant contained local excitatory feedforward connections and no learned lateral connectivity, with lateral communication restricted to the layer normalization operation. We reason that this model was capable of producing topographic organization in a way similar to the Self-Organizing Map (SOM) (34) and other algorithms applied to early visual cortex topographic organization (26, 28). Each of these algorithms implements a form of local cooperation alongside broader competition. Specifically, in the SOM, global competition is implemented by selecting a winning unit on each trial, and suppressing the responses of all other units, and local cooperation is mediated through Hebbian learning scaled by a Gaussian neighborhood around the winning unit. In ITN models including our feed-forward variant, the local excitatory feedforward connections implement a form of local cooperation, ensuring that neighboring units are positively correlated; the layer normalization operation then implements a global competition by attempting to convert the distribution of pre-activations to a standard normal distribution, which leads to sparser activity following rectification (the degree of which can be controlled by each unit’s bias term), and ensures that units represent different aspects of the feature space. Thus, layer normalization implements both competition and interactivity that, when combined with the local representational cooperation induced by local excitatory feedforward connections, leads to a smooth topographic organization whereby the unit feature tuning is systematically more similar for nearby units than for farther units. In recurrent ITN models, the learned lateral connections can adapt this competition and interactivity, allowing for increased performance (Figure 5).

Despite some conceptual similarities, there are several advantages to variants of the ITN architecture relative to SOMs and other previous topographic mapping algorithms. The first is that ITNs are naturally hierarchical, allowing for multiple interacting levels of topographically organized representations, rather than assuming a single feature space to be arranged in a single topographic map. This allows them to explain the presence of multiple domain-selective regions arranged in a stream from earlier to later parts of IT (1, 3, 66, 67). Second, and relatedly, the connectivity constraints of the ITN can be incorporated into generic task-optimized neural networks, without requiring separate Hebbian updates to topographically organize the feature space following development of the feature space as in the SOM. Lastly, the ITN framework is extremely flexible, allowing for future research to examine different encoders, different IT architectures and topologies, and different task training environments and readout mechanisms. This makes the ITN an attractive approach for future research examining topographic organization in the visual system.

One limitation of our current work is that it only addresses the topographic organization of high-level representations, since the connectivity constraints were not applied within the convolutional layers of the encoder network that was used to model early and mid-level vision. In deep learning architectures, convolutions are a crucial aspect of achieving good task performance, whereas local connectivity suffers from a relative lack of inductive bias, thereby requiring more parameters and longer training time to learn similar features at different visual field locations. However, this is a particular challenge for the ITN framework that also points to a critical limitation of convolutional architectures as a model of the brain. Attempting to model topographic organization in convolutional layers over both retinotopic location and stimulus features—well known organizing principles of early visual cortex—would necessitate that each channel have potentially different connections with other channels across different retinotopic positions, violating the convolution. In the brain, feature tuning is not actually uniform across the visual field (68). Thus, relaxing the convolution assumption has merits for advancing visual computational neuroscience, and would enable more detailed connectivity-based topographic modeling of early and mid-level visual areas, an important line of work that deserves future attention. Fully connected visual “Transformer” layers using multiplicative attentional interactions (69, 70) may prove to be a promising architecture in which to examine topographic organization using the ITN framework, as these architectures have recently been shown to reach high performance without convolutions.

The work presented here makes important progress in modeling, both quantitatively and qualitatively, the factors underlying visual cortical development throughout the visual hierarchy. Here, we focused on constraints local to the IT circuit. However, a currently unexplored question in our framework is why and how regions emerge in consistent locations across individuals of a given primate species (3, 17, 46, 71, 72). We hypothesize that modeling long-range connectivity-based constraints with regions external to IT (e.g., 44, 45) (see also 73), along with adapting the ITN architecture to contain two hemispheres, will give rise to reliable localization of model cortical areas based on their connectivity with upstream and downstream areas. In particular, the retinotopic organization of upstream early visual cortical areas is thought to encourage foveally-biased cortex to support face representations, and peripherally-biased cortex to support scene representations (45, 74). Moreover, innate connectivity biases with downstream nonvisual areas is thought to play a further role in shaping the global organization of domain-selective areas in IT (45, 75–79). These biases, such as left-hemispheric language biases, and other more fine-grained patterning of connections with domain-selective downstream areas (i.e., socially-responsive areas for faces, memory areas for scenes, motor areas for manipulable objects) should be explored in future work to better understand IT organization both within and between hemispheres. Based on previous work (43–45), we fully expect graded connectivity to bias the resulting locations of domain-selective regions. However, based on the results here, we argue that such long-range connectivity is not a necessary condition for topographic domain-selectivity; rather, the pressure for low wiring cost solutions to hierarchical visual computation within IT appears to be sufficient to drive such organization.

## Materials and Methods

### The Interactive Topographic Network

Here, we introduce the Interactive Topographic Network (ITN), a framework for computational modeling of high-level visual cortex, under specific biological constraints and in the service of specific task demands. ITN operates according to a set of principles which build upon previous work (45), and can be divided into two components: an *encoder* that approximates early visual cortex, and *interactive topography* (IT) layers that approximate inferotemporal cortex. The goal of the encoder is to extract general visual features which describe the visual world along dimensions that support a broad range of downstream readout tasks. However, our main modeling focus is on IT, which consists of a series of pairs of recurrent layers that are subject to biological constraints. For computational simplicity, such constraints are not modeled in the encoder, although future work that incorporated similar constraints could be used to model topographic organization throughout the visual hierarchy.

### Encoder architecture and training

We used a ResNet-50 (80) encoder to allow the ITN to extract deep and predictive features of the trained inputs. The encoder is pre-trained on equal sized subsets of faces, objects, and scenes from the VGGFace2 (81), ImageNet (82), and Places365 (83) datasets, respectively, matched in terms of total training images. We reused the same subsets of faces and objects as in (84), and an additional scene domain was constructed to match the other two domains in total images. An initial learning rate of 0.01 was used, and this learning rate was decayed 5 times by a factor of 10 upon plateau of the validation error; after the 5th learning rate decay, the next validation error plateau determined the end of training. Stochastic gradient descent with momentum (*ρ* = 0.9) and *l*2 weight decay (*λ* = 0.0001) was used, with batch size of 256 on a single GPU.

### Recurrent neural network formulation of IT

Our model of IT extends the standard discrete-time Recurrent Neural Network (RNN) formulation common in computational neuroscience (e.g., 85). We begin with the continuous-time dynamics of units in an RNN layer, where ***x***^(*a*)^ is the vector of pre-activation activities in area *a* of IT, ***r***^(*a*)^ is the vector of post-activation activities in area *a*, ***b***^(*a*)^ is the vector of baseline activities in area *a, τ* is the scalar neuronal time constant, and *W* ^(*a,b*)^ is the matrix of weights from area *a* to area *b*:

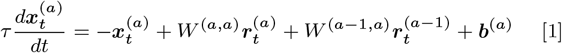

where the activation function 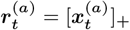 is positive rectification, also called a Rectified Linear Unit (ReLU). Applying the Euler method to integrate this first-order ordinary differential equation, with time step size Δ*t*, and substituting 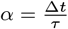, yields the discrete time update:

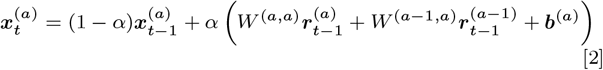

Note that this formulation differs from the standard machine learning implementation of RNNs, which can be derived as a special case where Δ*t* = *τ* or *α* = 1, in which the time constant is set such that the previous activity of a neuron decays exactly to zero in the time between updates, such that it can be set to 0 in the update equation.

When training models with separate excitatory and inhibitory units, we noted that training could be extremely unstable and required some mechanism for achieving stability. To this end, we adopted layer normalization (86), without the trainable scaling parameter that is sometimes used (see 86, for more details). Where *µ*(***x***) is the mean of ***x***, and *σ*(***x***) is the standard deviation of ***x***, and ***b*** is the learned bias term (moved outside of the layer normalization), the layer-normalized activities are given as:

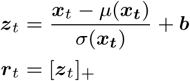

Incorporating layer normalization into our update equation yields the final update equation:

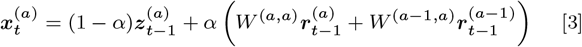

### Extending the standard RNN framework with biological constraints

Here, we outline the major biological constraints implemented in this work.

### Spatial organization

An essential aspect of an ITN model is that each IT layer has a spatial organization. We chose to model layers as square grids, with each layer of the hierarchy of equal size (typically, a grid size length of 32, corresponding to a layer of 1024 units). We normalize the coordinates to lie in the range [0,1]. Each unit thus has a unique (*x, y*) coordinate which will be used to determine the distance-dependent network topology. In general, the specific choices about map spatial arrangement are not critical to the predictions of the model, but they can potentially be manipulated in certain ways in the service of other theoretical goals.

### Spatial connectivity costs

We impose distance-dependent constraints on connectivity through a cost on longer connections throughout training. This basic formulation of the loss was introduced by (43) as a way to induce spatially organized task specialization, and was shown to do so in a simple neural network model trained on small-scale tasks. To our knowledge, no other research has examined this loss in modern deep learning architectures trained on natural images. We use a simple modification of the original loss function, using the squared Euclidean distance 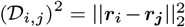 (in place of 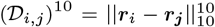 distance (43)). By using the squared distance, we penalize longer connections disproportionally compared to shorter connections. The spatial loss on connections between areas *a* and 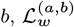, is given by:

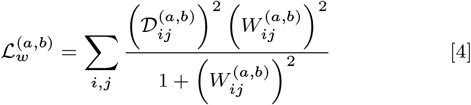

The total spatial loss is the sum of the area-to-area spatial losses 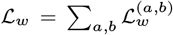 and is added to the task-based loss as ℒ = ℒ_*t*_ + *λ*_*w*_ ℒ_*w*_, on which gradient descent is performed.

### Connection noise

To approximate axon-specific variability in instantaneous firing rate (87), We apply multiplicative noise on the individual connections between neurons which is uniform over distance and layers. In practice, we find that connection noise helps to regularize the activations in the network, encouraging a more distributed representation. Noise is sampled independently from a Gaussian distribution 𝒩 centered at 0 with variance *σ*^2^ at each time step of each trial, and is squashed by a sigmoidal function 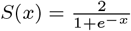, ensuring that the sign of each weight is not changed and each magnitude does not change by more than 100%. Thus, the noisy weight matrix 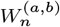 from area *a* to area *b* on a given trial and time step is:

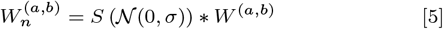

### Sign-based restrictions on neuronal connectivity

As has been discussed in prior computational work (85), standard neural networks gloss over a key detail of neuronal morphology—that single neurons obey Dale’s Law, whereby all of their outgoing connections are either excitatory or inhibitory (ignoring modulatory neurons and other special, rare cases). We employ this principle within our framework by replacing the single sheet of unconstrained neurons with parallel sheets of excitatory (E) and inhibitory (I) neurons. We follow the method of (85) to enforce the sign of connectivity in our neuronal populations. The second sign-based restriction we implement is that between-area interactions are carried out predominantly by excitatory pyramidal neurons. Thus, we restrict between-area feed-forward connectivity to originate from the excitatory neurons only. Both E and I neurons receive feedforward inputs.

### Task demands

Task-driven computational models learn representations that better account for neural responses in visual cortex than models which are designed by hand (24, 25, 49). More recent work has shown that a supervised version of task-driven learning is not essential, with semi-supervised contrastive learning algorithms performing very close to the supervised state-of-the-art in neural prediction (88). For simplicity and as a first step, we use supervised learning: Given a set of categories, the network is tasked with classifying images into one of these categories. The cross-entropy loss is used as an objective function, optimized using stochastic gradient descent with weight decay and momentum. We expect our general findings to hold for any learning algorithm that places comparable demands on high-level representation learning.

Visual systems often have to perform multiple tasks, where the specific organization of tasks is not known ahead of time. Therefore, a pre-specified task segregation (c.f. 89) is not possible. By specifying only a domain-general architecture, and optimizing to performance over multiple tasks, ITN models can discover the task-specific organization that maximizes task performance in the context of its task-general architectural constraints. Here, we simulate the requirement of performing three somewhat different visual tasks from common inputs, using common resources. The first task is face identification, for which we use VGGFace2 (81). The second task is object and animal recognition (hereafter just referred to as objects), for which we use the ImageNet dataset (82). The third task is scene recognition, for which we use the Place365 dataset (83). These three tasks constitute three separate “domains,” each containing several categories which the network must learn to discriminate between. However, the network has no prior knowledge of the separability of these domains, and they are fully interleaved during training.

### IT architecture and training

The main ITN model consists of 3 IT layers (pIT, cIT, aIT) with separate E and I populations, and feedforward connections sent only by E units. To facilitate training many models with fewer computational demands, the model is trained using a fixed pre-trained ResNet-50 encoder on smaller subsets of faces, objects, and scenes. Specifically, we created image subsets equal to the size of the popular CIFAR-100 dataset but at higher image resolution, containing 100 categories each with 500 training images and 100 validation images, resized to 112×112 pixels. Thus, the combined dataset contained 300 categories with 150,000 training images and 30,000 validation images. The same learning rate schedule as used for training the encoder was used. Stochastic gradient descent with momentum (*ρ* = 0.9) was used, with batch size of 1024 on a single GPU. We used spatial regularization with *λ*_*w*_ = 0.05, without additional weight decay on IT connections.

### IT model variants

To better understand the factors that contribute to the development of topographic organization, we examine a variety of IT model variants containing different subsets of implemented constraints. Some of these models do not use separate populations of E and I units, but still restrict feedforward connectivity to be excitatory. In this case, we simply restrict the feedforward weights to be positive, despite the same neuron having both positive and negative lateral connections. In another variant, we remove learned lateral connections entirely. This model is trained for a single time step, and the only recurrent computation is that of a single pass of layer normalization. Lastly, we explore a range of spatial regularization strengths.

### Analyses of trained models

After training, the responses in IT layers are probed to investigate emergent task specialization and its topographic organization. We use three main approaches.

### Mass univariate analyses

The first analytic approach is the simple mass-univariate approach, in which each unit is analyzed separately for its mean response to each stimulus domain (objects, faces, scenes), using untrained validation images from the same categories used in training. In addition to computing the mean response to each domain, we compute *selectivity*, a ubiquitous metric used in neuroscience, to analyze how responsive a unit is to one domain compared to all others. We compare the responses of each domain versus the others using a two-tailed *t*-test, and given the test statistic *t*, the significance value *p* of the test, and the sign of the test statistic *s* = *sign*(*t*), we compute the selectivity as −*s* log(*p*).

### Searchlight decoding analysis

The second analysis approach we use is the multivariate searchlight analysis commonly used in fMRI (51), in which a pool of units are selected in a (circular) spatial window around each unit, and the accuracy for discriminating between different categories (e.g., hammer vs. screw-driver) in each domain (e.g., objects) is computed using the activations of only that pool of units; the mean accuracy value is assigned to the center unit, and the process is repeated for all units.

### Lesion analysis

To assess the causal role of certain units in the performance of specific tasks, we adopt a lesioning approach in which the activities of lesioned units are set to 0 throughout perception. This effectively removes them from processing, allowing the network’s dynamics to unfold independently of these units. The effect of a lesion is measured by computing the accuracy following the lesion and relating that to the baseline accuracy.

The first type of lesion we perform is a spatial or *focal* lesion in which a circular neighborhood of size *p* ∗ *n* units is selected, where *p* is the fraction of units selected and *n* is the total number of units in the area where the lesion is performed. The lesion is centered on a unit *u*_*i,j*_ either randomly or according to the peak of a specific metric such as selectivity. In the main analyses, we attempt to lesion spatial neighborhoods corresponding to regions of high domain selectivity. To do so, we take the selectivity map, perform spatial smoothing, and select the unit *u* of peak smoothed selectivity. We then systematically vary *p* while keeping *u* fixed to assess the causal role of increasingly large regions centered on the peak of smoothed selectivity.

The second type of lesion sorts units according to a given *selectivity* metric irrespective of their spatial location. In this analysis, the *p* ∗ *n* most selective units are chosen for a given lesion. This is done separately for the selectivity of each domain, as in the focal lesions. When the topography is smooth and the regions approximately circular, the selectivity-ordered and focal lesions yield similar results. However, to the extent that the topography is not perfectly smooth or circular, the selectivity-ordered lesion may knock-out a more relevant set of units for a given task.

### Distance-dependent response correlation

We calculate the correlations of the responses of all pairs of units as a function of distance between them. Response correlation is computed for a given time step over a large number of images, either from all domains, or from each domain separately. We additionally compute a scalar metric of this analysis by taking the Spearman correlation of response correlation and distance. This metric can be easily visualized over many time steps, layers, cell types, models, etc.

### Analyzing spatial costs of trained networks

To understand the wiring cost of certain trained models, we analyze the spatial cost of a network, as given by Equation 4, as a function of architectural parameters such as the spatial regularization strength *λ*_*w*_. In one analysis, we analyze only the feedforward spatial cost, which simply requires summing spatial costs over pairs of areas *a* and *b* where *a*! = *b*. Similarly, to analyze only the recurrent spatial cost, we can sum spatial cost over pairs of areas *a* and *b* where *a* = *b*.

### Unweighted spatial cost of sparsified networks

While wiring cost in an artificial neural network should depend to some extent on the strength of connections—stronger connections may require greater myelination, and strong connections in an artificial neural network may correspond to a larger number of connections in a biological neural network—there is another notion of wiring cost whereby it depends only on whether or not two neurons are connected. This notion of wiring costs has been commonly applied to the study of cortical areal layout and early visual cortical maps (e.g. 29, 58, 59, 90). Moreover, the analysis of binary connectivity in thresholded networks is also common in graph-theoretic analysis of brain data (91). To analyze this notion of wiring costs, we pruned our trained models to a desired connection sparsity level *s*, setting to 0 the *n*∗*m*∗*s* connections with the smallest magnitude, where *n* and *m* are the number of units in areas *a* and *b*. Sparsity was enforced globally within IT and from IT to readout, rather than individually for each set of connections. We then analyzed an unweighted wiring cost 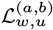 that computes the mean of squared Euclidean distance values between connected units *i* and *j* in areas *a* and *b*, given that (*a, b*) are in the set of connected areas *C*:

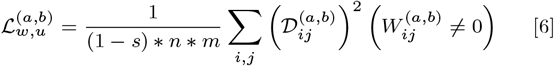

## ACKNOWLEDGMENTS

The authors thank Michael J. Tarr, Leila Wehbe, Vlad Ayzenberg, Jacob Prince and the VisCog research group at Carnegie Mellon University for helpful discussions and comments on this work. N.M.B. thanks Rosemary A. Cowell for early discussions that helped to conceive of the work.

