## Supporting Information for "A connectivity-constrained computational account of topographic organization in primate high-level visual cortex"

Nicholas M. Blauch[1,2] Marlene Behrmann[2,3] David C. Plaut[2,3]
[1]Program in Neural Computation [2]Neuroscience Institute [3]Department of Psychology
Carnegie Mellon University
{blauch,behrmann,plaut}@cmu.edu

July 8, 2021

#### 1 Final model performance

We measured overall recognition performance of the main ITN model throughout training. The results are plotted for training and validation images as a function of training epoch in Supplementary Figure S1. Additionally, we plotted the final accuracy for each domain as a function of processing time step in the model.

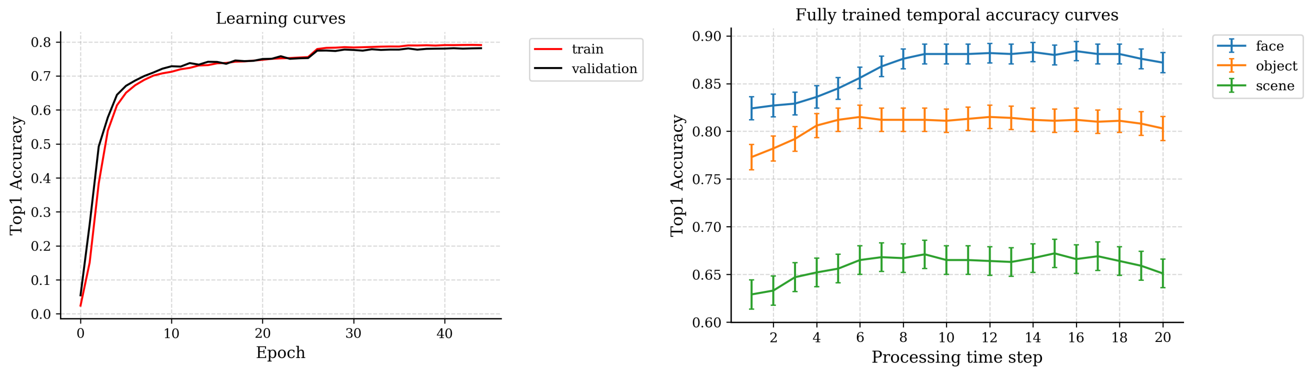


**Supplementary Figure S1:** Performance of the main model. **A.** Overall accuracy throughout the training of IT using a pre-trained encoder. **B.** Final accuracy for each domain, plotted over model processing time steps.

#### 2 A more complete assessment of domain-selective functional organization

In the main paper, for clarity, we plotted lesion deficits at two specific lesion sizes. Here, for completeness, we plot both raw accuracy values and lesion deficits for a large range of lesions, shown in Supplementary Figure S2.

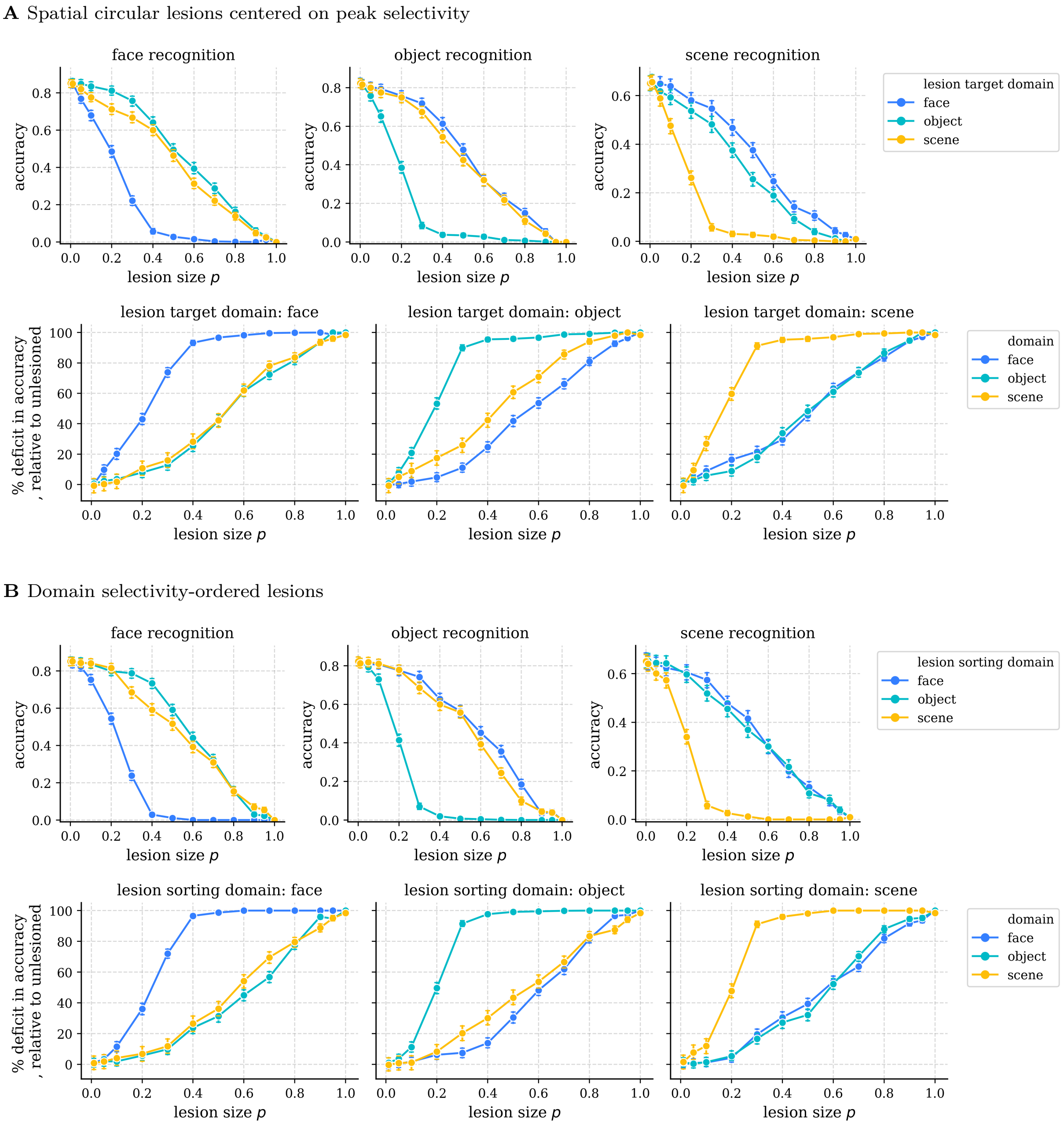


**Supplementary Figure S2:** Lesion results in the ITN model. **A.** Damage from various sizes of circular topographic lesions centered on the peak of smoothed selectivity for each domain. **B.** Damage from various sizes of non-topographic lesions chosen by sorting units according to their selectivity for each domain. As the selectivity is highly topographic, the lesion masks in the topographic vs. selectivity-ordered lesion plots are very similar; however, for selective regions that are non-circular (such as that seen for objects), the selectivity-ordered lesion provides a more precise way of assaying the selective region.

To better understand the degree of representational competition and cooperation, we compare searchlight accuracy and mean readout maps across domains. The results are plotted in Supplementary Figure S3 as scatter plots over units, colored by their selectivity between the two plotted domains, for all pairs of domain. The results corroborate our earlier findings, demonstrating a strong but graded degree of specialization. In particular, we note the large degree of correlation between searchlight accuracies for objects and for scenes. This result is somewhat surprising given the relatively specialized effects of lesions, demonstrating that object and scene information may co-mingle but still be read out from different largely different sets of units in order to optimize task performance.

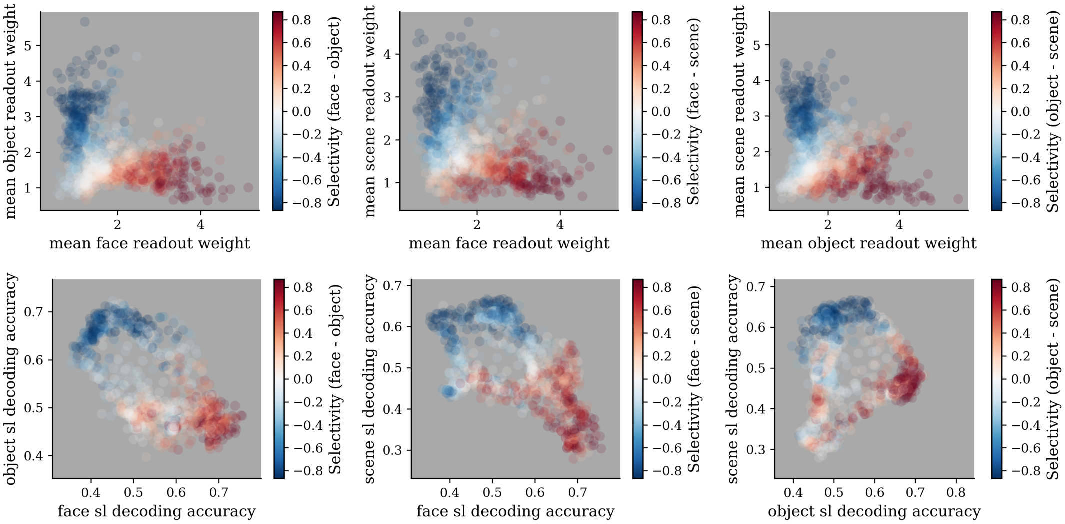


**Supplementary Figure S3:** Graded competition and cooperation between domains in aIT. Top: readout weights into each domain, bottom: searchlight accuracy for decoding within each domain.

How consistent is the topographic responsiveness of categories within a domain? Within a given domain-selective area, are the weaker responses to non-preferred domains roughly uniform and weak across categories, or do some categories elicit notably strong responses? To answer this question, we plotted the mean aIT response and readout weights of 10 randomly selected categories from each domain, with the contour of significant smoothed domain-selectivity ($p<0.001$; smoothing performed as averaging selectivity over the 5% nearest units). These results are shown in Supplementary Figure S4.

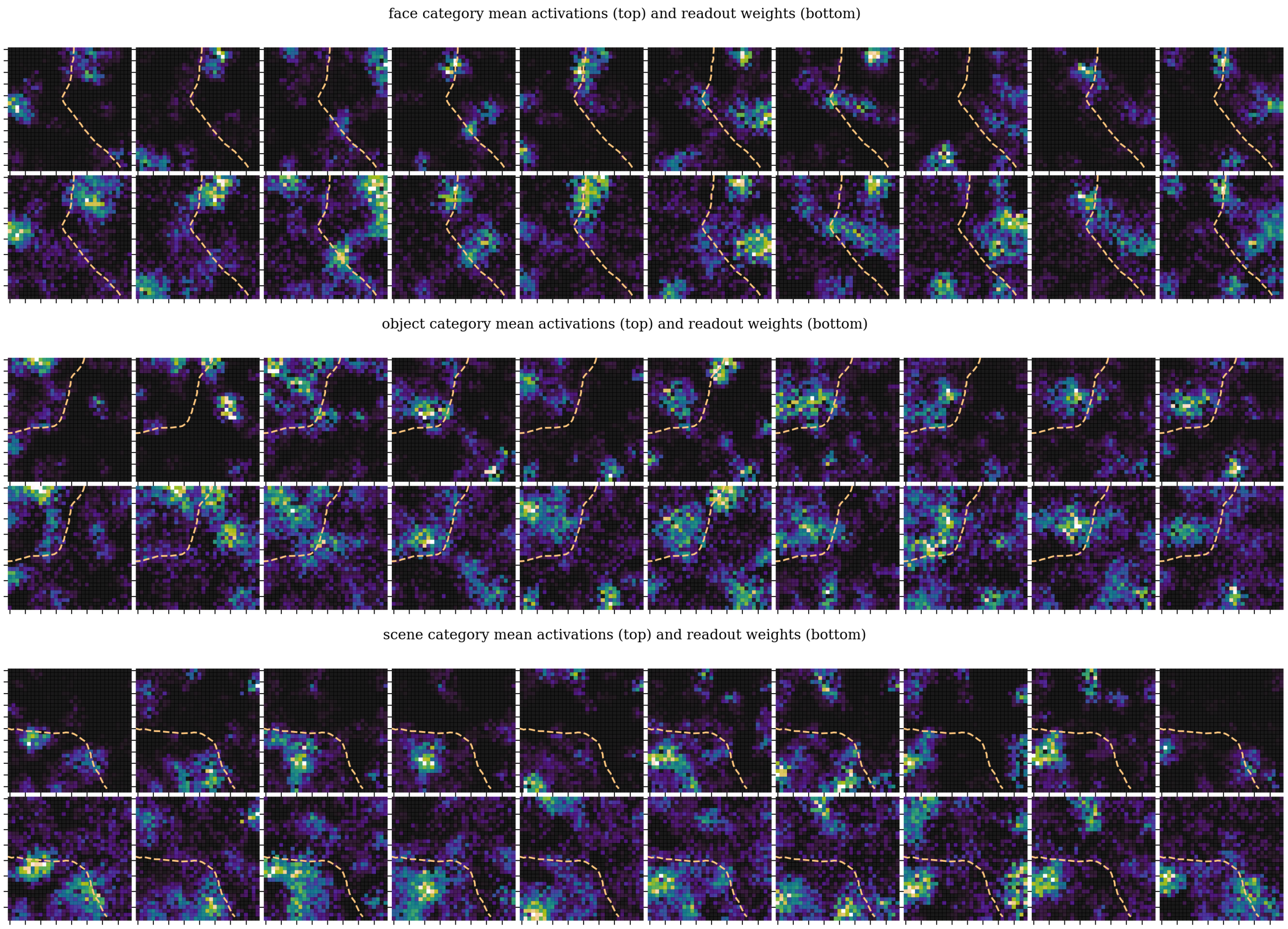


**Supplementary Figure S4:** Category-level mean responses and readout weights for 10 example categories from each domain. For each domain, the top row shows mean category-level responses in aIT, and the bottom row shows the readout weights for the corresponding categories. Dashed yellow lines indicate the contour of statistically significant smoothed domain selectivity for the plotted category's domain ($p<0.001$; smoothing performed as averaging selectivity over the 5% nearest units).

### 3 Detailed results for the simplified feedforward model

In the main paper, we presented a simplified "feedforward" model that utilizes excitatory feedforward connectivity and layer normalization. In Supplementary Figure S5, we plot a larger range of results for this model, including domain selectivity, within-domain information, lesion deficits, and generic topography.

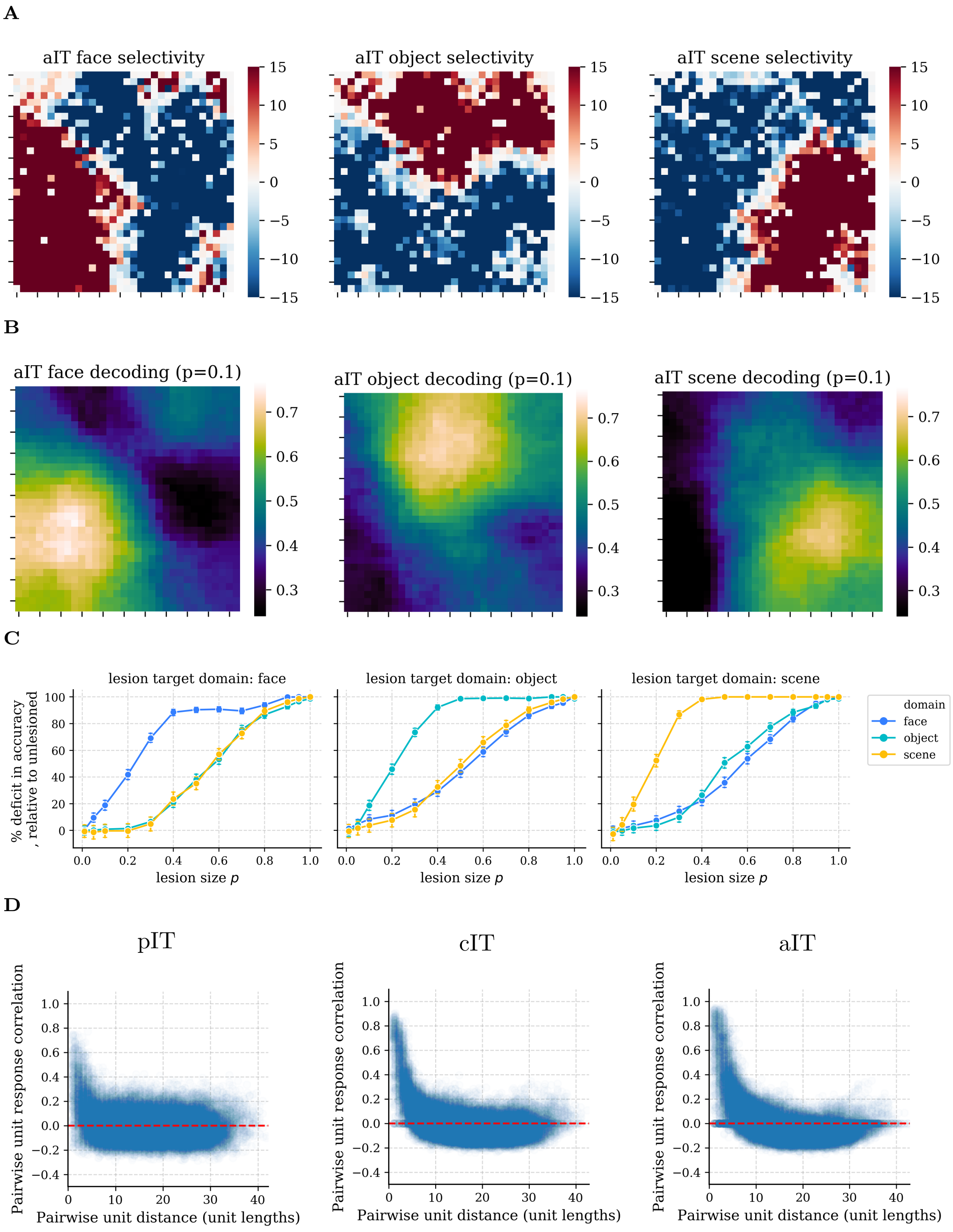


**Supplementary Figure S5:** A simplified model containing local, excitatory feedforward connectivity and global lateral divisive inhibition (layer normalization) exhibits domain-level and generic topographic organization very similar to the more biologically-detailed model. See Figures **??** and **??** for details on the plots.

### 4 Detailed results for an intermediate model, with strictly excitatory feedforward connections but no separation of E and I

In the main paper, we presented a simplified "recurrent" model that utilizes excitatory feedforward connectivity, layer normalization, and recurrent lateral connectivity. In Supplementary Figure S6, we plot a larger range of results for this model, including domain selectivity, within-domain information, lesion deficits, and generic topography.

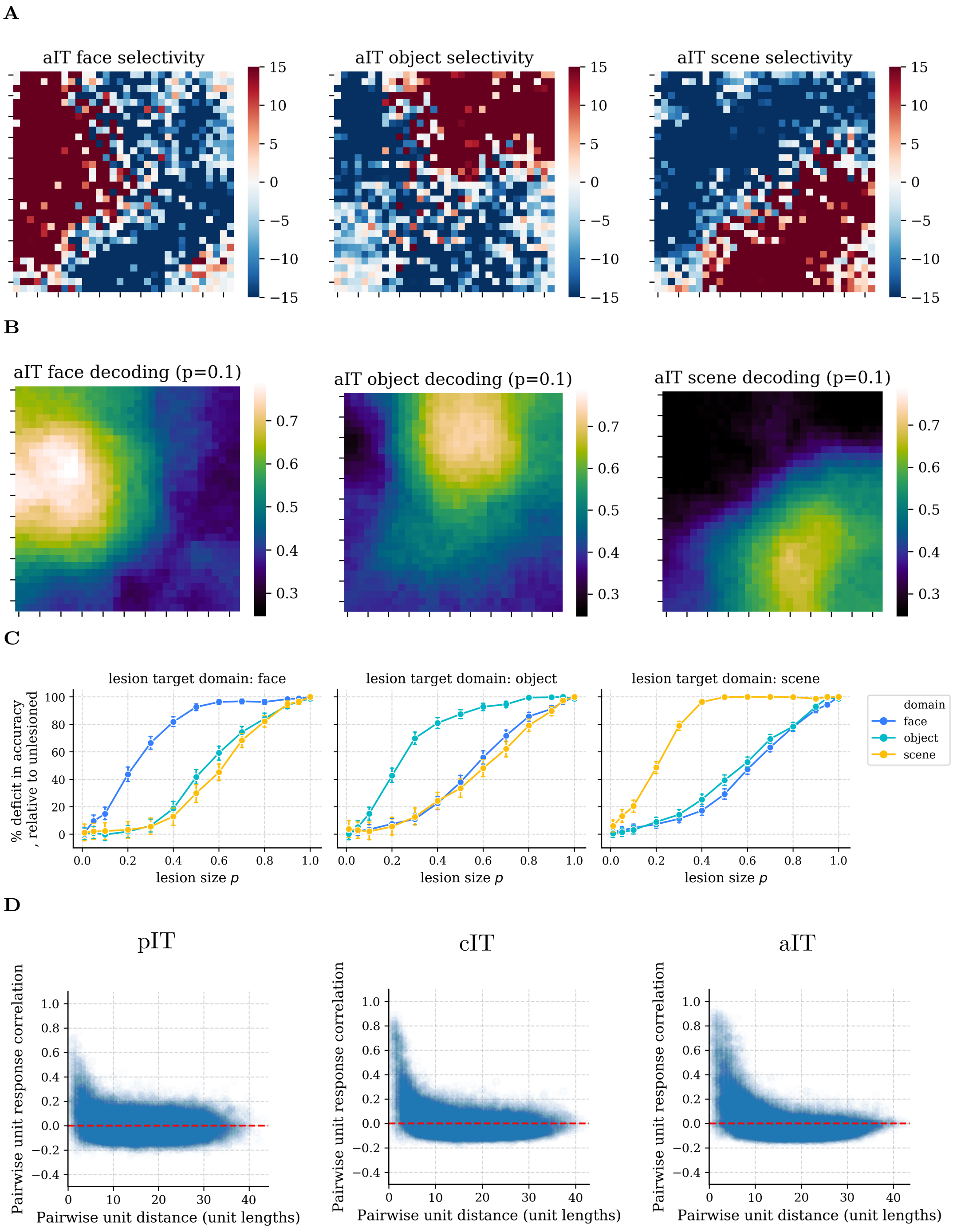


**Supplementary Figure S6:** A simplified model containing local, excitatory feedforward connectivity, local lateral connectivity (one sheet of units without sign constraints), and global lateral divisive inhibition (layer normalization) exhibits domain-level and generic topographic organization very similar to the more biologically-detailed model.

### 5 Clustering results for the face domain

In Supplementary Figure S7, we show a reduced analysis that preserves privacy of face images, plotting only the cluster centroids and quantification of the attributes of biological sex and whether the modal hair color is blonde (inferred from a sample of 8 images per identity) to illustrate that within-domain clustering is seen for faces along with the other domains shown in the main paper. As with the other domains, further analysis of other possible attributes (e.g. ethnicity, age) is possible, but is beyond the scope of this work.

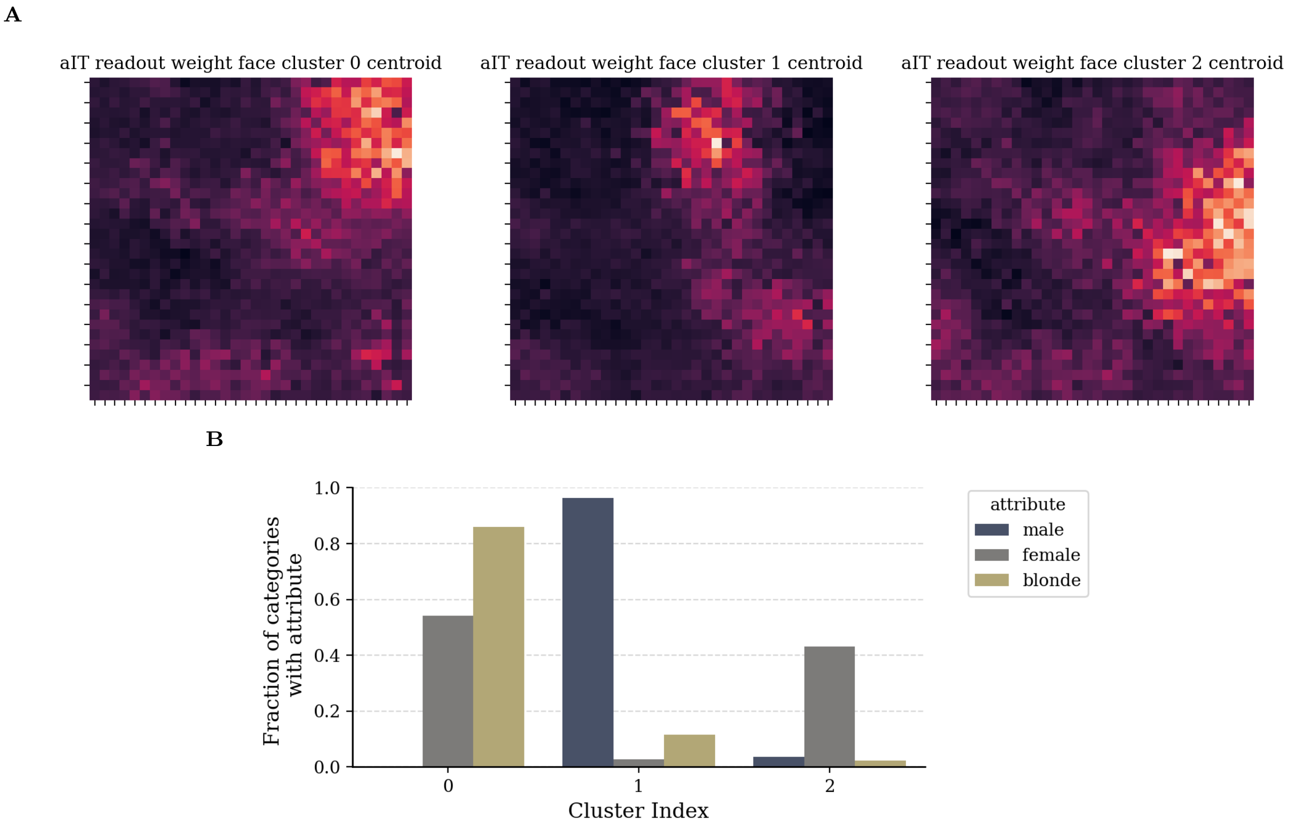


**Supplementary Figure S7:** Clustering face category readout weights from aIT. $k = 3$ clusters were used in a K-means++ clustering algorithm that clustered the vectors of aIT readout weights over all 100 face categories. **A.** The centroids of the clusters were then reshaped into the 2D aIT coordinates and visualized as heat maps. **B.** Quantification of biological sex over the members of each cluster, revealing quantitative characteristics of each cluster.

### 6 Clustering for the scene domain

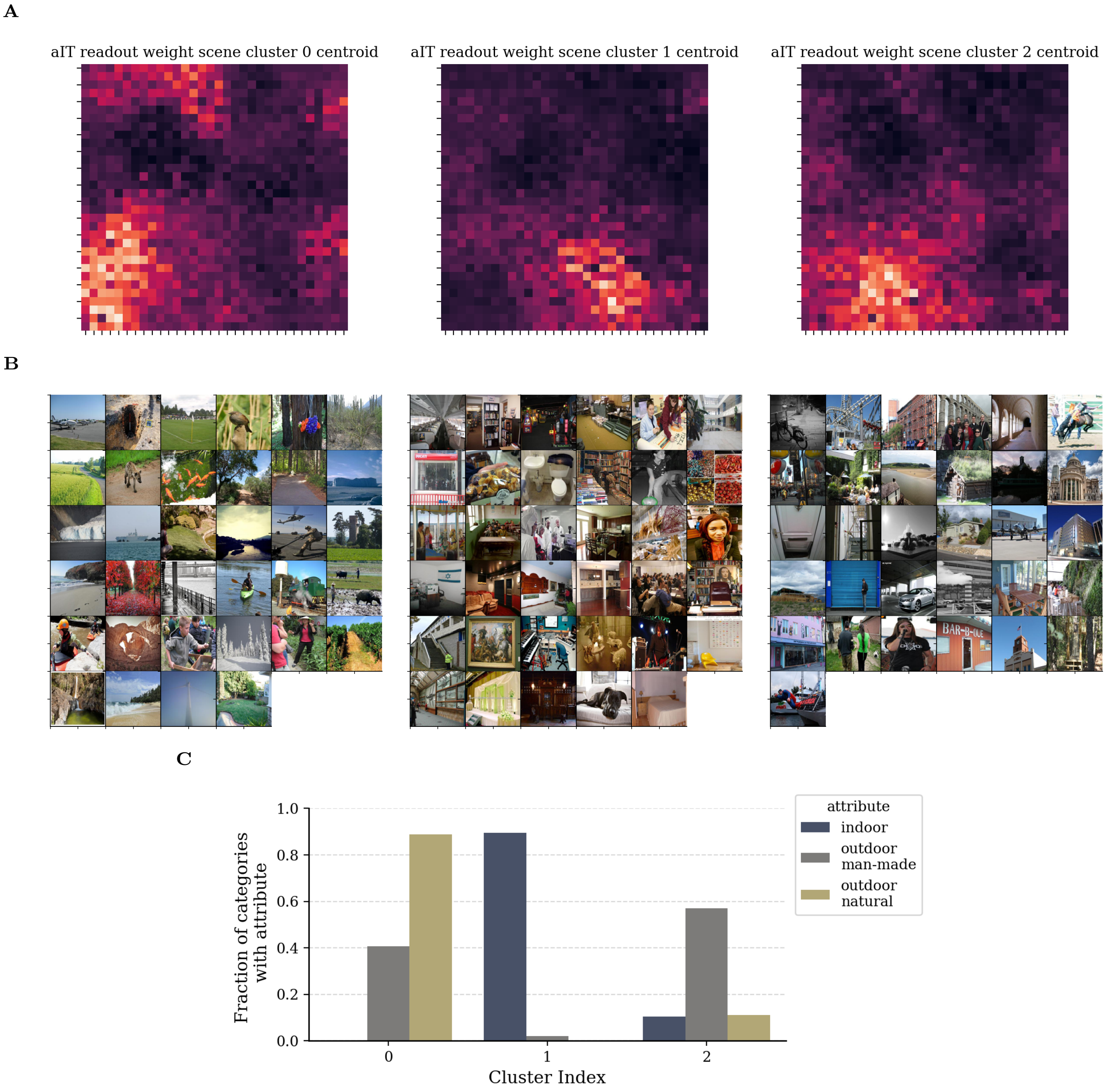


**Supplementary Figure S8:** Clustering scene category readout weights from aIT. $k = 3$ clusters were used in a K-means++ clustering algorithm that clustered the vectors of aIT readout weights over all 100 scene categories. **A.** The centroids of the clusters were then reshaped into the 2D aIT coordinates and visualized as heat maps. **B.** Cluster category members. **C.** Scene attribute quantification.

### 7 The impact of visual domain experience on emergent domain-level topography

To assess the dependence of emergent topography on visual experience, we trained three additional ITN models with the E/I-EFF-local-RNN architecture, each trained on only one domain (objects, faces, or scenes), with all other parameters identical. The encoder was pre-trained with the same domain images used to train the encoder in main model, removing the images from the other domains. Similarly, IT was trained with the same domain images used to train IT in the main model, removing the images from the other domains. Domain-level topography and generic distance-dependent response correlations are plotted in Supplementary Figure S9. Somewhat smooth domain responses, along with an inverse relationship between pairwise unit distance and response correlation, are seen in each model. However, the model trained on all domains produces the cleanest global topographic organization, followed by object- and scene-trained models. The face-trained model appears to demonstrate the most specific topographic organization, biases towards faces, which makes sense in light of the more specialized data present in cropped (but naturalistic) face pictures compared to naturalistic photographs of objects and scenes.

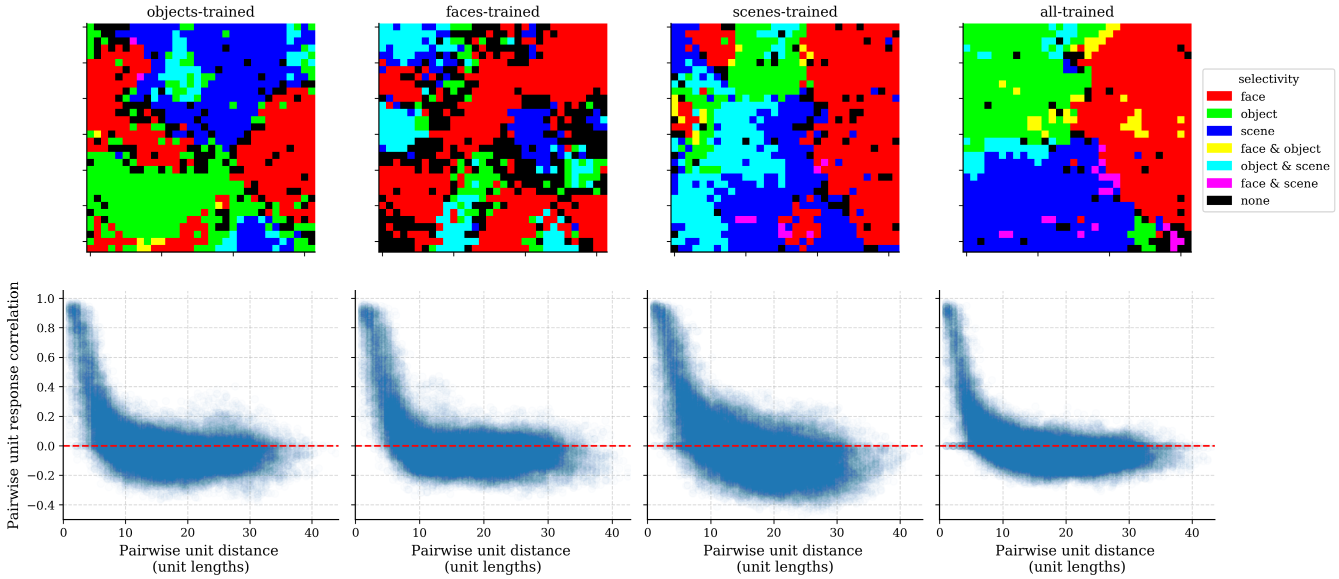


**Supplementary Figure S9:** Topographic organization under different visual training experiences. The main model trained on objects, faces, and scenes is compared with models trained on each of the domains separately. As in the main model, a pre-trained encoder was used to learn the topography in the IT layers; in each case, the specific training environment was applied to both the encoder and IT components, using a larger dataset for pre-training the encoder.

### 8 Robustness of results

To verify the robustness of our results, we ran multiple versions of the main ITN model using different random seeds controlling the random weight initialization and training stimulus presentation order, both of which can in theory effect the emergent functional organization. We plot these results in Supplementary Figure S10.

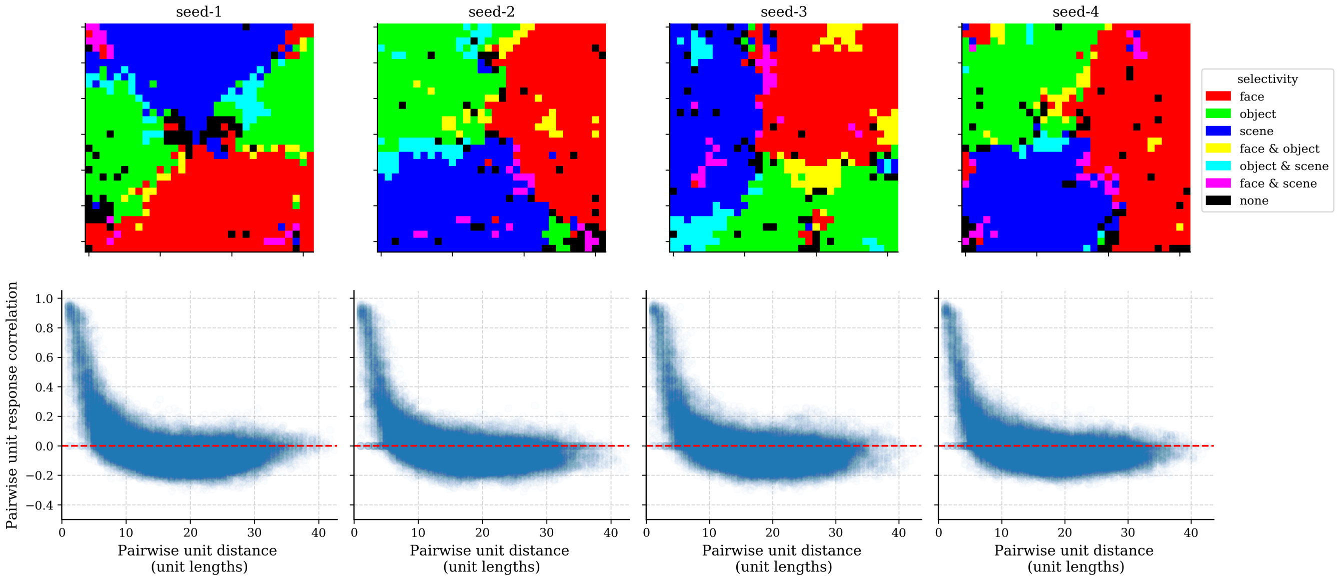


**Supplementary Figure S10:** Robustness of results. Top: domain-selective topography in aIT. Bottom: generic distance-dependent response correlations in aIT.
